# DNA barcoding of passerine birds at an ornithological crossroad reveals significant East-West genetic lineage divergence

**DOI:** 10.1101/2024.12.12.628090

**Authors:** Sahar Javaheri Tehrani, Elham Rezazadeh, Niloofar Alaei Kakhki, Leila Nourani, Vali Ebadi, Sahar Karimi, Mojtaba Karami, Fatemeh Ashouri, Asaad Sarshar, Toni I. Gossmann, Mansour Aliabadian

## Abstract

Exploring genetic diversity is essential for precise species delimitation, especially within taxonomically complex groups like passerine birds. Traditional morphological methods often fail to resolve species boundaries; however, DNA barcoding, particularly through the mitochondrial cytochrome c oxidase subunit I (*COI*) gene, provides a powerful alternative method for accurate species identification. This study establishes a comprehensive DNA barcode library for Iranian passerine birds, analyzing 537 *COI* sequences from 94 species across 23 families and 53 genera. We observed a pronounced barcode gap, with average intraspecific divergence at 0.4% and interspecific divergence at 18.6%. Notable intraspecific variation emerged in the Persian nuthatch (*Sitta tephronota*) and the Lesser whitethroat (*Curruca curruca*), while the goldfinch (*Carduelis carduelis*) showed limited genetic differentiation despite marked morphological distinctions. Phylogenetic analysis revealed significant east-west genetic splits in *C. curruca* and *S. tephronota*, reflecting geographic and zoogeographic boundaries of Iran. These findings demonstrate the effectiveness of DNA barcoding in elucidating biogeographic patterns, emphasizing the key role of Iran as an ornithological crossroads for avian biodiversity. Moreover, our results suggest that much of the genetic variation in the *COI* gene arises from synonymous mutations, highlighting the role of purifying selection in shaping mtDNA diversity across species.

## Introduction

Genetic diversity is a fundamental aspect of biodiversity, representing the variety of genetic information within and among species (Nonić and Šijačić-Nikolić 2021). This genetic diversity plays a critical role in evolutionary processes such as natural selection and adaptation, driving speciation and shaping the phylogenetic relationships among organisms (Huang et al. 2016; Nonić and Šijačić-Nikolić 2021). Accurate assessment of genetic diversity is also fundamental for species delimitation, particularly in cryptic or morphologically similar species, where traditional taxonomic methods may fall short (Hebert et al. 2003; Lohman et al. 2009; Bilgin et al. 2016). Different genetic markers offer varying effectiveness in studying genetic diversity and it is recommended to use fast changing molecular markers (i.e., coding vs. noncoding DNA) for closely related species (Abdel-Mawgood 2012). DNA barcoding is a transformative technique in biodiversity research, allowing for the precise identification and differentiation of species through genetic markers. This method utilizes a standardized, short segment of the mitochondrial gene cytochrome c oxidase I (*COI*), typically a 648-base pair region, to serve as an internal species tag for species delimitation (Arida et al. 2021; Zhang and Bu 2022). This species delimitation relies on DNA barcoding gap, which refers to the difference between mean intraspecific and interspecific genetic distances (Antil et al. 2023) and beyond taxonomy (Chac and Thinh 2023), this approach has also been employed in studies of biogeography, ecology and biological conservation (Gostel and Kress 2022 b; Wu et al. 2023). Moreover, genetic diversity levels in mitochondrial DNA are influenced by various factors, primarily mutation rate, selection, and effective population size (Clark et al. 2023). Understanding the selective forces acting on *COI* sequences provides valuable insights into species evolutionary histories and adaptive responses, with important implications for biodiversity conservation strategies and for tracking ecological changes over time (Matzen da Silva et al. 2011).

Birds represent one of the most extensively studied animal groups in DNA barcoding projects (Hebert et al. 2004), achieving species-level identification accuracy ranging from 93 to 99% (Colihueque et al. 2021). This high level of accuracy demonstrates the effectiveness of DNA barcoding in discriminating among avian species (Colihueque et al. 2021). While significant research on avian diversity has been concentrated in Europe and North America (Grant et al. 2021), DNA barcoding efforts have also extended to various other regions, such as the Nearctic (Hebert et al. 2004), Neotropics (Kerr et al. 2009a; Kerr et al. 2009b; Tavares et al. 2011), South Korea (Yoo et al. 2006), eastern Palearctic (Kerr et al. 2009b), Scandinavia (Johnsen et al. 2010), Indomalaya (Lohman et al. 2009; Lohman et al. 2010), and Australasia (Patel et al. 2010). Consequently, DNA barcodes are currently available for approximately 41% of bird species worldwide, encompassing about 4,300 species from 37 of the 39 recognized avian orders (Colihueque et al. 2021). Despite the expansion of DNA barcode reference databases in species diversity and geographic coverage, (Gostel and Kress 2022 a; Cheng et al. 2023) many regions remain underrepresented, resulting in a notable geographic bias in barcoded species representation (Colihueque et al. 2021).This gap highlights an urgent need for more comprehensive sampling in these poorly documented areas (Gostel and Kress 2022 a). Furthermore, obtaining the required permits for specimen collection and transporting samples across national borders is often particularly complex, especially for birds (Lijtmaer et al. 2012), which adds to the challenges addressing biodiversity gaps.

Iran is recognized as a globally significant biodiversity hotspot, characterized by its remarkable species richness and high levels of endemism (Aliabadian et al. 2005; Aliabadian et al. 2007; Noori et al. 2024; Rezazadeh et al. 2024). This diversity can be largely attributed to the country’s geographic complexity, steep climatic gradients, and pronounced landscape heterogeneity (Noori et al. 2024). Additionally, Iran’s unique geographic position serves as a zoogeographical transition zone where several major biogeographical realms—the Palearctic (both eastern and western), Oriental, and Afrotropical—intersect (Noori et al. 2024). This strategic positioning not only establishes Iran as an ornithological crossroads but also contributes to the notable presence of sister bird species within the country (Aliabadian et al. 2005; Aliabadian et al. 2007). However, despite constituting a significantly more important hotspot for diversity, the population structure and genetic diversity of the passerine taxa— representing the most species rich clade of birds—remain inadequately explored within the country. To address this gap, we have conducted an extensive sampling effort to generate a comprehensive DNA barcoding library of Iranian Passerine birds.

Our main objectives are (i) to evaluate genetic variation in *COI* among passerine birds in Iran—a region characterized by numerous contact zones between passerine species (Aliabadian *et al*., 2007)—to provide new insights into the efficacy of *COI*-based DNA barcoding; (ii) to identify potential cryptic species; and (iii) to investigate the impact of natural selection on mitochondrial *COI* sequences.

## Material & methods

### Taxon sampling

The study area covers the northeastern and western regions of Iran (Fig. S1). We examined 537 individuals representing 94 species from all these regions, with 75 species of these taxa (80 % of whole samples) represented by more than two individuals. Birds were captured using mist nets, identified, and then released following the collection of feather and blood samples. Blood samples were collected from the brachial vein of each bird following standard protocols and preserved in Queens’s buffer (Seutin et al. 1991). No birds were harmed during the capture, handling, and blood collection process. For the taxonomy of species, we used the IOC World Bird List v14.2 (Gill et al. 2024). The complete list of sampled specimens including information about geographical location, voucher number and, access numbers is provided in (Table S1).

### Laboratory procedures

DNA was extracted from blood and feather samples using a standard salt extraction method (Brookefieldand and Burke 1992), following overnight incubation at 40°C in an extraction buffer containing 2% sodium dodecyl sulfate (SDS) and 0.5 mg/ml proteinase K. Additionally, 30 µl DTT was added during the initial incubation step for feather extraction. The *COI* gene was selected as the molecular marker of choice, as it is recognized as the standard DNA barcoding marker by the Consortium for the Barcode of Life (CBOL) and the All Birds Barcoding Initiative (ABBI) (Hebert et al. 2004). Primer pairs and a locus-specific annealing temperature that have been used to amplify this gene region are shown in (Table 1). Total PCR reaction volumes were 25 μl, containing 12.5 μl Taq DNA Polymerase Master Mix RED (Ampliqon), 1 μl of each primer with a concentration of 10 μM, 3 μl DNA, and 7.5 μl ddH2O. PCR products were examined on 2% agarose gels to confirm the successful amplification of the target fragments. The purified PCR products for all specimens were sequenced by Macrogen Inc (Seoul, South Korea).

**Table 1.**
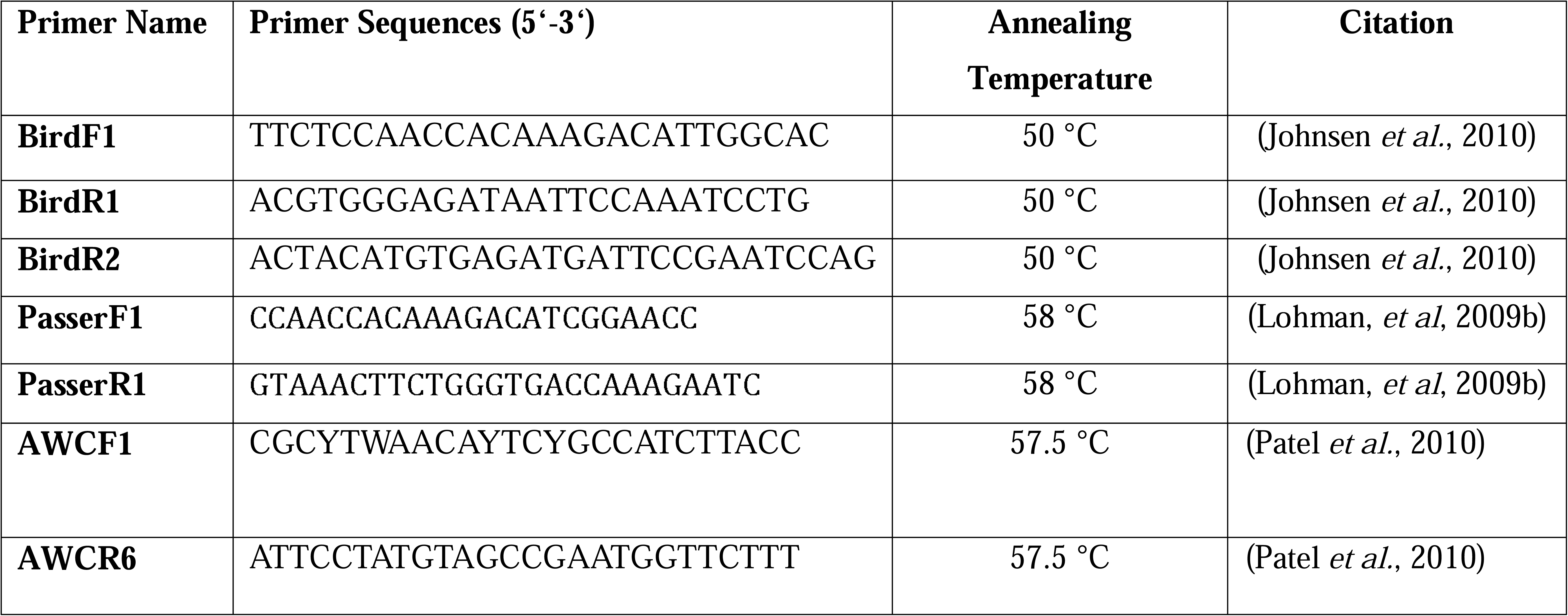
Primer pairs that have been successfully used to obtain bird barcodes. This table includes forward and reverse primer names, primer sequences, annealing temperature, and citation.

### Data analysis

Sequences were aligned and edited in BIOEDIT v7.0.1 (Hall 1999). Intraspecific and interspecific distances were calculated using Kimura 2-parameter (K_2_P) pairwise genetic distances in MEGA v6.0 (Tamura et al. 2013). Average intraspecific distances were determined for species with at least two specimens using MEGA. The (K_2_P) model was employed for all sequence comparisons, as it is considered the most effective metric for evaluating closely related taxa (Nei and Kumar 2000). The best-fit model was estimated using JMODELTEST v2.1. (Darriba and Posada 2014) based on the Bayesian Information Criterion (BIC). The best model was then used to construct a phylogenetic tree using MRBAYES v3.2.0 (Ronquist and Huelsenbeck 2003) to provide a general graphic representation of the pattern of divergence between all species. The phylogenetic tree was rooted with one representative of Galliformes (*Gallus gallus*). A species was considered by DNA barcode if: a) it was monophyletic (i.e., the species formed a single cluster) and b) it did not share a barcode with any other species.

Consequently, high intraspecific genetic distances in the *COI* gene are frequently utilized to predict cryptic or potentially new species. In our dataset, this pattern is observed in two species showing elevated genetic distances: the Persian Nuthatch *Sitta tephronota* Sharpe, 1872 and the Lesser Whitethroat *Curruca curruca* Linnaeus, 1758 (Table S2). Additionally, our dataset includes the Goldfinch *Carduelis carduelis* Linnaeus, 1758, recognized as a complex species with phenotypically distinct hybridizing subspecies groups (Haffer 1977) and ongoing uncertainties regarding its species status of certain subspecies (Abdilzadeh et al. 2023), warranting further investigation.

For these three taxa, a Neighbor-Joining (NJ) tree was constructed using K_2_P distances in MEGA. This analysis incorporated the sequences obtained in this study and those deposited in BOLD (https://www.barcodinglife.org) from Iran. Furthermore, a haplotype network was implemented in POPART v1.7 (Leigh et al. 2015) to visualize the relationships among haplotypes. Pairwise K_2_P distances between populations were estimated with the program MEGA, and pairwise F_ST_ values were calculated with DNASP v5.1 (Librado and Rozas 2009)

### Coding DNA Genetic diversity analysis

Based on the *COI* sequence fragments and the subsequent global alignment we obtained genetic diversity at 0-fold and 4-fold sites for all species. For this, we used the vertebrate mitochondrial genetic code with MEGA. We then used the Tajimas_d package from the bfx suite (https://py-bfx.readthedocs.io/en/latest/) to calculate nucleotide diversity for each species for 0-fold and 4-fold sites, respectively. We excluded species with zero diversity for either 4-fold or 0-fold sites. One can quantify effective population by dividing genetic diversity with the mutation rate per generation (π = 2Ne * µ, where Ne equals the effective population size, µ, the mutations per generation and π is the observed pairwise differences in a population genetic sample). Because we have limited knowledge of mitochondrial gene specific mutation rates for all passerine birds and only rough estimates for generation time, we cannot directly estimate effective population sizes. However, here we use genetic diversity at silent sites as a proxy for effective population size, which is not unreasonable because we restrict our analysis to passerine birds, a taxonomic group with supposedly little variation in mutation rate and generation time.

## Results

### COI sequence variation

A total of 537 sequences for 94 passerine bird species were generated and uploaded to the BOLD database (both publicly available, Table S1) which belong to 53 different genera and 23 different families of Passeriformes order. The average number of sequences per species was six (1-32). The mean intraspecific K_2_P distance was 0.04 % (range from 0-3.61 %) and the mean interspecific distance was 18.6 % (Fig. 1).

**Figure 1.**
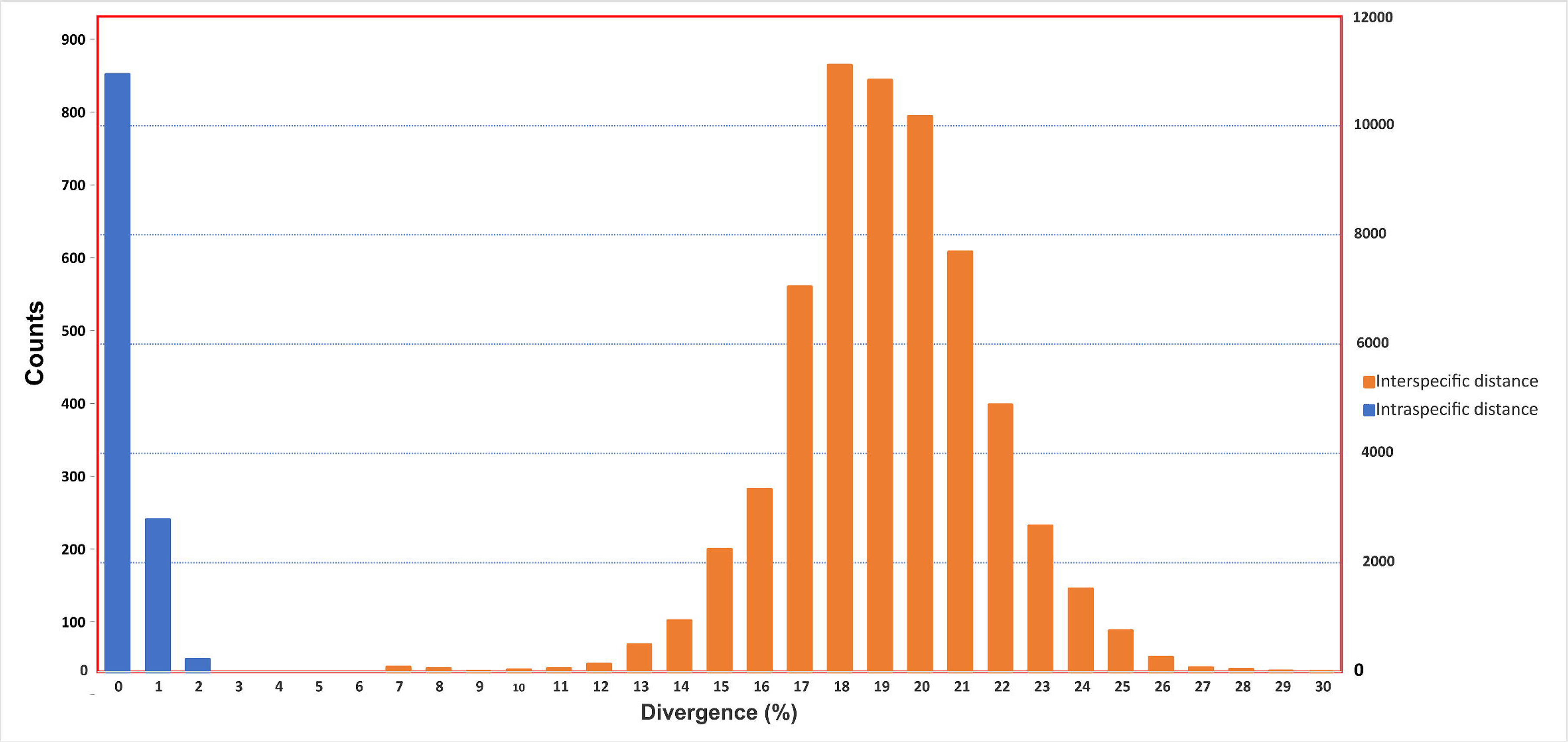

We reconstructed a phylogenetic tree with a Bayesian approach using all the specimens (537 sequences, Fig. S2). In the obtained phylogenetic tree, most of the nodes are well resolved and strongly supported with posterior probability (PP) more than 95%, as indicated by bold lines in the tree. Of the 23 families analysed, 20 (excluding Muscicapidae, Emberizidae and Fringilidae) were found to be monophyletic. Similarly, 51 of the 53 genera (excluding *Emberiza*, and *Luscinia*) appeared monophyletic. Furthermore, all species formed well-supported (PP > 95%) monophyletic groups except, *Lanius collurio* which displayed a paraphyletic pattern (Fig. S2).

Although most species showed relatively low intraspecific distances, two species showed high intraspecific K_2_P distances clearly above the threshold of 2 to 3% sequence divergence in our data set including *Sitta tephronota* (2.3%) and *Curruca curruca* (3.61%). Conversely, our analyses revealed low intraspecific genetic distance in the *Carduelis carduelis* complex (0.43%), which contrasts with the significant morphological differentiation observed (Table S2). In order to illustrate the basic pattern in these three taxa, their results were presented in detail in (Figs 2-4 respectively).

**Figure 2.**
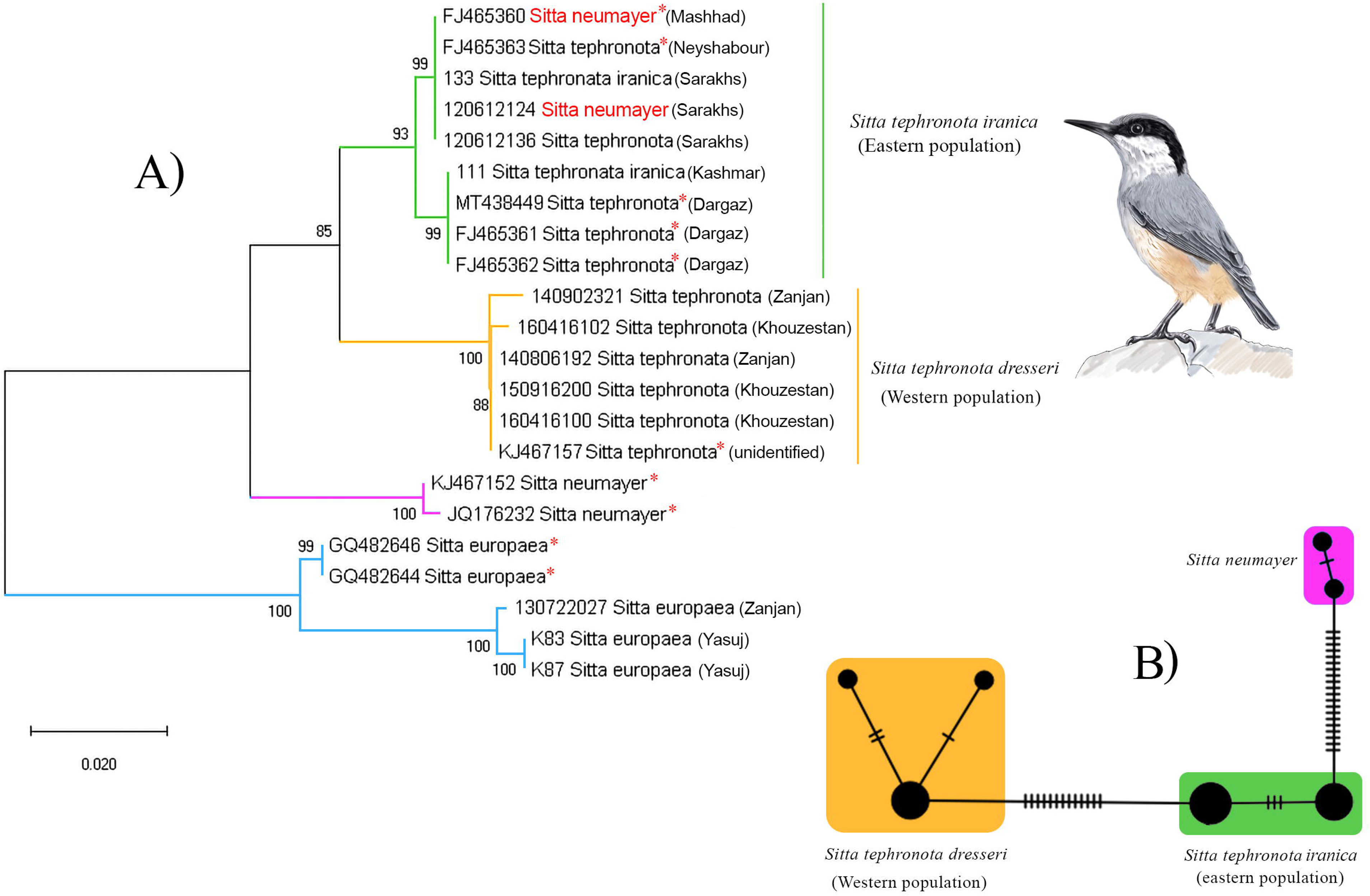

### Deep and shallow intraspecific divergences

*Sitta tephronota* includes three subspecies in Iran, i.e., *S. t. dresseri* (Zagros Mts. in SE Turkey to N Iraq and W Iran), *S. t. obscura* (NE Turkey to the Caucasus and Iran) and *S. t. iranica* (NE Iran and S Turkmenistan). For this species, we analysed six samples from the western population (*S. t. dresseri*) and seven samples from the eastern population (*S. t. iranica*). Two eastern-western major clades with high support were identified through both NJ and Bayesian analysis (Fig. 2 and Fig. S2). All *S. t. dresseri* samples formed a strongly supported clade, which is the sister group to another well-supported clade containing samples from the eastern population (Fig. 2A). This differentiation of haplotypes into two clades in the NJ tree was also mirrored by the presence of two haplotype groups in the *S. tephronota* network. In the haplotype network two haplotype groups were separated by 13 base pairs (Fig. 2B). The pairwise F_ST_ and genetic distances between eastern-western populations of *S. tephronota* are 0.91 and 4.1% respectively (Table 2 and Table S3). Furthermore, two individuals, which are recognized as *Sitta neumayer* in BLAST, are clustered with one of the eastern subclades of *S. tephronota*.

**Table 2.**
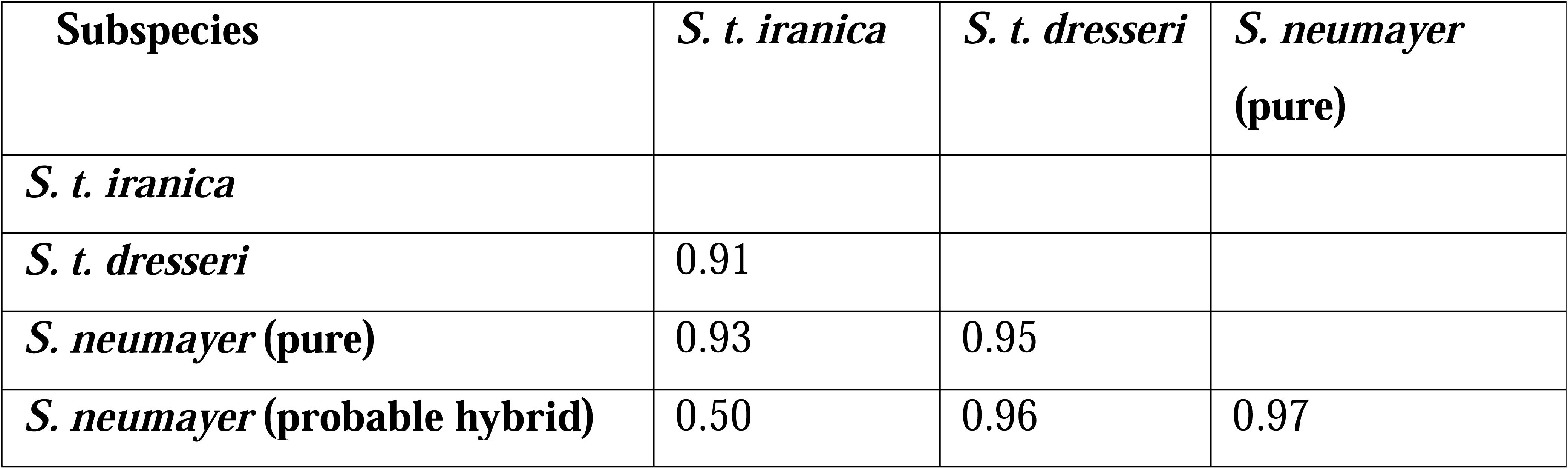
Pairwise F_ST_ values between the studied subspecies and species of *S. tephronota* and *S. neumayer* estimated from the mitochondrial data.

*Curruca curruca* is proposed to have three breeding subspecies in Iran, including *minula*, *althaea*, and *curruca* and one non breeding subspecies *halimodendri* Sarudny, 1911. For this species, two major eastern-western clades with high support were identified through both NJ and Bayesian analysis (Fig. 3 and Fig. S2). Furthermore, the eastern clade (including samples from Khorasan province) is divided into two well-supported subclades. One subclades represent *althaea* and another new eastern subclade aligns with the distribution range of *halimodendri* and/or *minula* in northeast Iran (Olsson et al. 2013; Abdilzadeh et al. 2023) Additionally, another major, well-supported western clade contains samples from the western part of Iran, within the distribution range of *C. c. curruca* (Fig. 3A). In the haplotype network, these two eastern-western populations were separated from each other by 21 base pairs (Fig. 3B). The pairwise F_ST_ and genetic distances between *C. curruca* subspecies from eastern and western populations in Iran are as follow: *curruca*/*althaea* (F_ST_ = 0.99; K_2_P = 7.04%), *althaea*/new subclade (F_ST_ = 0.96; K_2_P = 1.20%) and new subclade/*curruca* (F_ST_ = 1.00; K_2_P = 6.99%) (Table 3 and Table S4 respectively).

**Figure 3.**
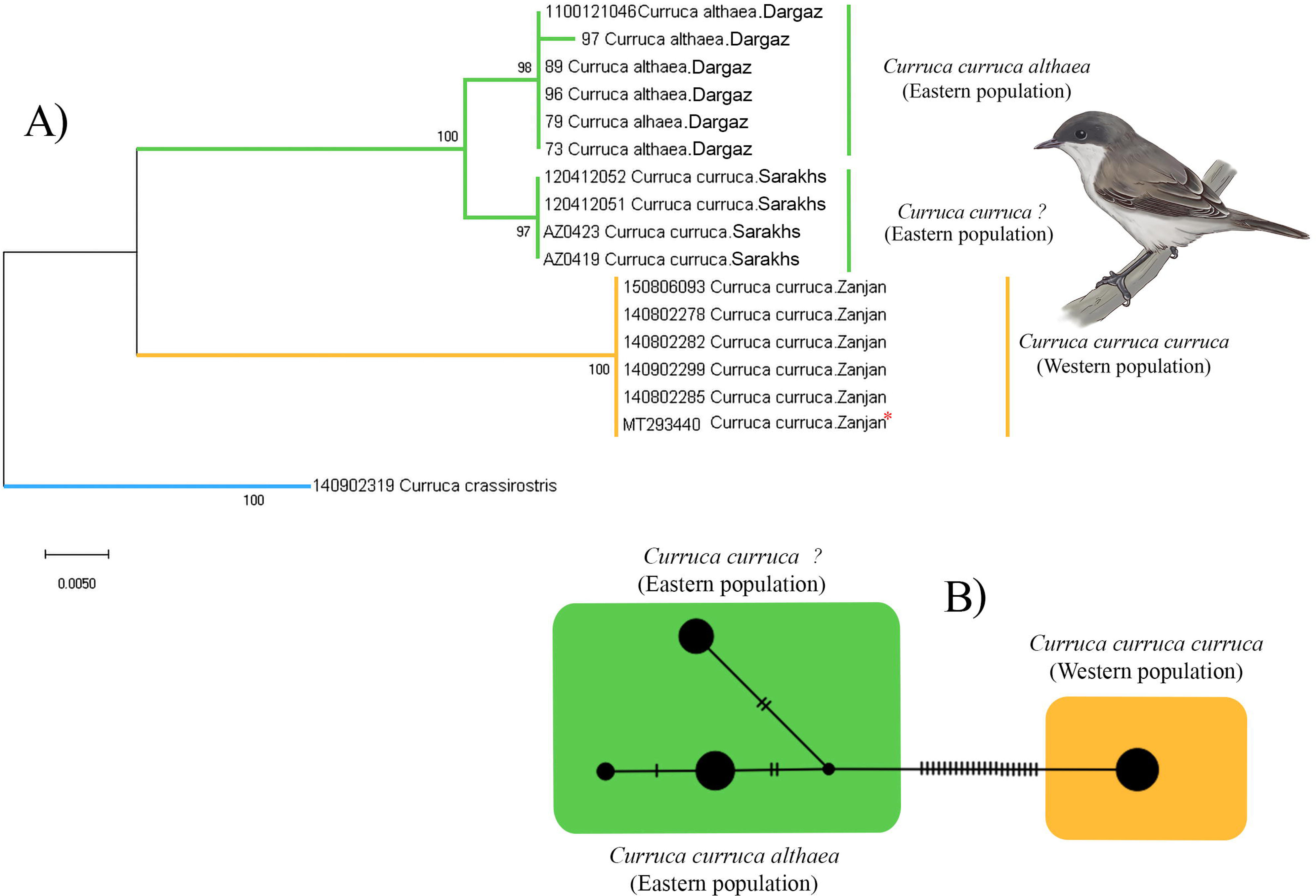

**Table 3.** Pairwise F_ST_ values between the studied subspecies and species of *Curruca curruca* estimated from the mitochondrial data.

| Subspecies | *C. c. curruca* (east.new subclade) | *C. c. curruca* (west) |
| --- | --- | --- |
| *C. c. curruca* (east. new subclade) |  |  |
| *C. c. curruca* (west) | 1.00 |  |
| *C. c. althaea* (east) | 0.960 | 0.992 |

*Carduelis carduelis* in Iran is represented by two subspecies groups: *caniceps* group in the east and *carduelis* group in the west (Haffer 1977). For this species, 18 different samples from the western population (*carduelis group*) and 9 samples from the eastern population (*caniceps* group) were analysed. These samples formed two main eastern-western clades; however, they received insufficient support (Fig. 4A). In the haplotype network, these two eastern-western populations were separated by two base pairs (Fig. 4B). Furthermore, one individual sampled from west of Iran (Yasuj), was located in eastern clade in both phylogeny and haplotype network. The pairwise F_ST_ and genetic distances between eastern-western populations of *C. carduelis* were 0.71 and 0.43% respectively (Table 4 and Table S5, respectively).

**Figure 4.**
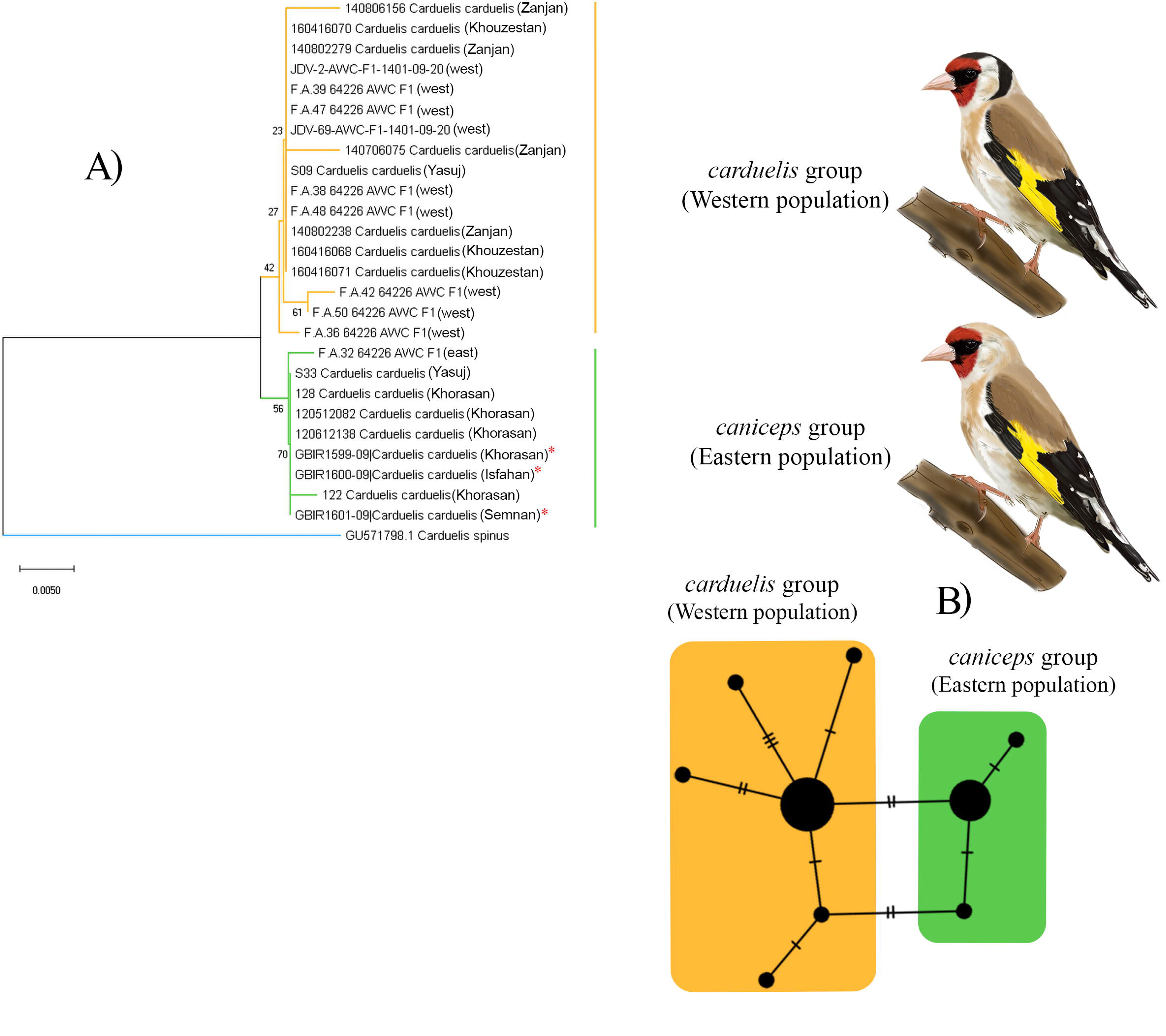

**Table 4.** Pairwise F_ST_ values between the studied subspecies and species of *Carduelis carduelis* estimated from the mitochondrial data.

| Subspecies group | *carduelis* group |
| --- | --- |
| *carduelis* group |  |
| *caniceps* group | 0.71925 |

### Patterns of COI gene variation

We quantified site-specific coding diversity (i.e., at 0-fold and 4-fold degenerate sites) of each species where we had multiple samples (Table S6). For subsequent analysis, we excluded all species with zero diversity at either of these site types and found for all remaining species that the logaritmized ratio of π 0-fold and π 4-fold is negatively correlated with π 4-fold (Fig. 5) which is consistent effective population size scaled effectiveness of selection (James et al. 2016)

**Figure 5.**
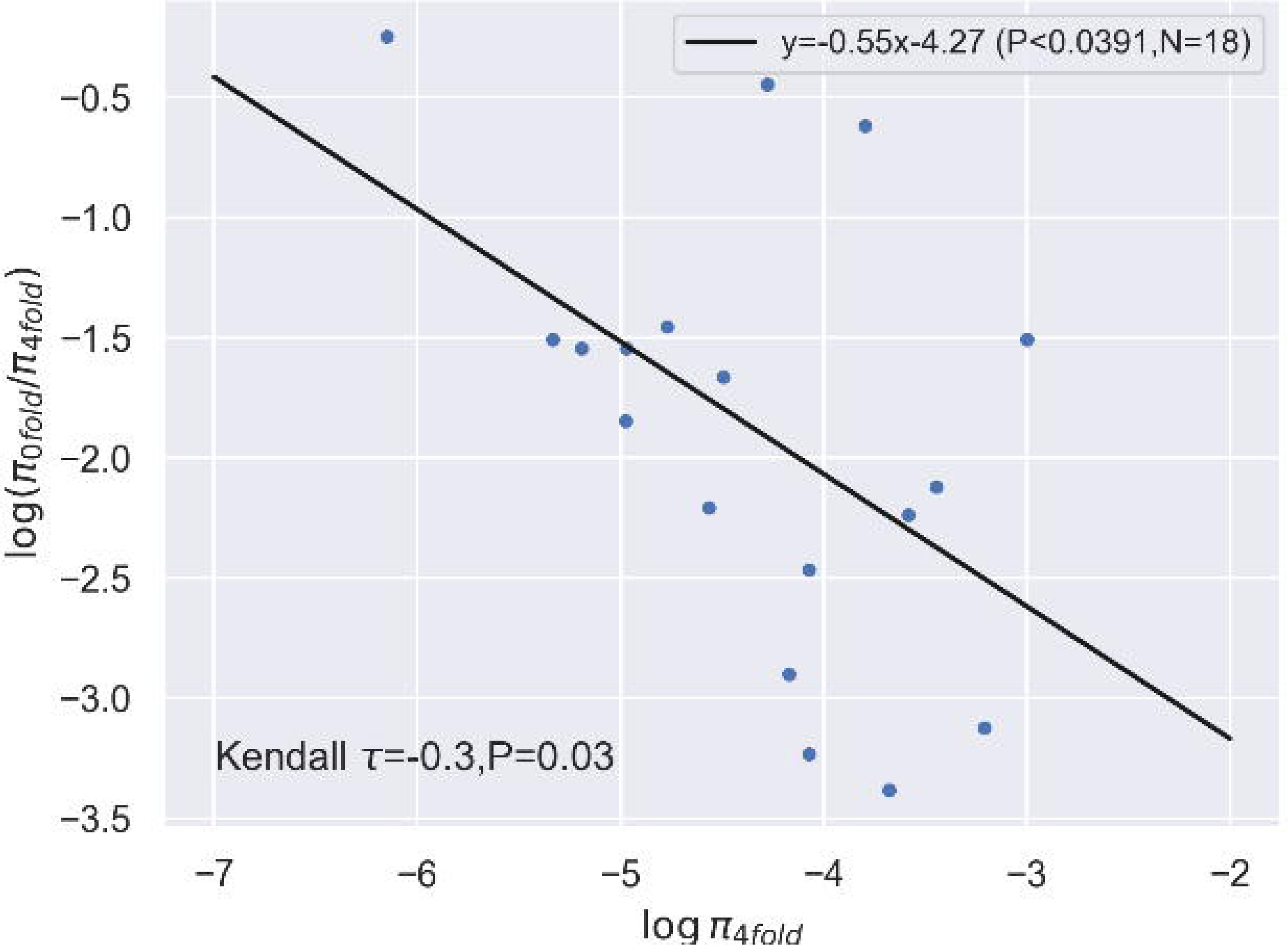

## Discussion

### DNA library for passerine bird species

Here, we provide a DNA barcode reference library for a substantial dataset of passerine birds in Iran, encompassing the identification of 94 distinct species. This study is the first comprehensive DNA barcoding-based assessment of Iranian passerine birds, featuring dense intraspecific sampling from both eastern and western regions. Our results again demonstrate that DNA barcoding is an effective tool for preliminary biodiversity assessments. No species shared sequences or had overlapping clades with any other species, and every passerine species had distinct *COI* sequences. The development of this DNA barcode library provides a valuable resource for the biodiversity of passerine birds in Iran and will facilitate future studies on the geographic variation and genetic diversity of passerine birds in that area.

Our results generally do not resolve phylogenetic relationships above the generic level for *Fringillidae*, *Emberizidae*, and *Muscicapidae*, at the generic level for *Luscinia* and *Emberizia* and, at the species level for *L. collurio*, all of which exhibited paraphyletic patterns in the phylogenetic analysis (Fig. S2). Therefore, using only *COI* alone cannot address higher-level taxonomic controversies in some cases. To further clarify the taxonomic uncertainties of higher passerine taxa, multiple nuclear markers using phylogenomic approaches (Zhao et al. 2023), potentially combined with detailed morphological comparisons, are required.

Below, we discuss our findings in relation to the genetic diversity of the three challenging taxa in Iran and the influence of the DNA barcoding in unrevealing biogeographic patterns.

### High intraspecific genetic distance and subtle morphological variation

Most DNA barcoding studies of birds frequently reveal genetically distinct yet morphologically cryptic species (Aliabadian et al. 2013; Saitoh et al. 2015; Bilgin et al. 2016). In our study, we identified two cryptic species, *S. tephronota* and *C. curruca,* which exhibits high intraspecific genetic variation while showing only subtle morphological differences. This highlights the challenges in distinguishing cryptic species based solely on traditional morphological methods, as these species may appear similar in morphology but harbor significant genetic divergence (Abdilzadeh et al. 2020). Our results support the subdivision of *S. tephronota* into two major well-supported east (*S. t. dresseri)* – west (*S. t. iranica)* clades (Fig. 2). This genetic split among Iranian populations of *S. tephronota* was first identified by Packert et al. (2020) in their efforts to clarify the phylogeny of the *Sitta* genus. They only have two samples from Iran, and they discovered a genetic divergence between a sample from eastern Iran (from Dargaz) and an unidentified sample referenced in Pasquet et al. (2014). However, they noted that it remains unclear whether the unidentified sample corresponds to *S. t. dresseri* or *S. t. iranica*. Based on our results, we conclude that this unidentified sample is likely representative of *S. t. dresseri*, as it clustered with other samples from western Iran.

Moreover, our finding showed that two individuals identified in BLAST as *S. neumayer* is located in the eastern clade of *S. tephronota* in both NJ tree and haplotype network. *S. tephronota* has an ecologically and morphologically similar congener, *S. neumayer*, which their distribution range overlap in eastern Turkey and Iran (Mohammadi et al. 2016). These two samples, which were collected from eastern Iran (Khorasan), might be hybrid samples, as it is primarily assumed that hybridization occurs between *S. tephronota* and *S. neumayer* in Iran (Haffer 1977). However, because mtDNA is typically maternally inherited, it is insufficient for identifying interspecific hybrids. Biparentally inherited nuclear gene marker is required to complement mtDNA data (Gruenthal and Burton 2005). Therefore, whether these two sister species meet without gene exchange or hybridized to a greater or lesser extent, additional sampling from their potential contact zone is needed. In addition, similar results were found by Elverici et al. (2021), when analyzing mitochondrial *ND2* and *ND3* gene sequences for *S. tephronota* and *S. neumayer*, and they indicated a reciprocal monophyly with no gene flow between birds in the Zagros Mountains and other populations. Furthermore, *S. europaea*, used as an outgroup for *Sitta* phylogenetic tree, exhibits genetic divergence between our samples from the western/northwestern population in Iran (*S. e. persica* and/or *S. e. rubiginosa*) and two other samples representative of the European haplotype. This finding aligns with previous results, which identified two additional Caspian mitochondrial lineages of *S. europaea* from Iran (Nazarizadeh et al. 2016).

*C. curruca* complex is an intricate model for studying cryptic speciation, presenting challenges in taxonomy due to conflicting morphological and genetic data (Abdilzadeh et al. 2020). Our analysis revealed a basal genetic split in this species between eastern (*althea)* and western (*curruca*) subspecies in Iran, supporting previous findings (Olsson et al. 2013; Abdilzadeh et al. 2020; Abdilzadeh et al. 2023).

Abdilzadeh et al. (2022, 2023) suggested that there is no geographical pattern between *caucasica*, *zagrossiensis* and *curruca* subspecies that all these three subspecies occurred in western Iran, and that all are synonyms due to their clustering. Our phylogenetic trees indicate two well-supported subclades within the eastern clade: one consisting of *althaea* birds, breeding in eastern Iran, and another sister subclade from lowland northeastern Iran (Sarakhs). It is unclear if this subclade represents *minula* or *halimodendri*.

Although, three subspecies *minula*, *althaea*, and *curruca* are believed to breed in Iran (Sarudny 1911), but records of *minula* from the eastern arid lowland areas are lacking aligning with Olsson et al. (2013). Our phylogenetic tree positions the new subclades as sister to *althaea,* conflicting with Olsson et al. (2013), *minula* is considered a sister taxon with *curruca*. Additionally, most previous authors have stated that there is no evidence to suggest that *minula* breeds outside of China (Olsson et al. 2013).

This evidence corroborate the assumption that our new subclade may not align with *minula*. *halimodendri* is another suggested taxon, which is assumed to have distribution in northeastern Iran (Olsson et al. 2013). Three *halimodendri* samples deposited in NCBI were all collected out of the breeding season from southeast Iran (Abdilzadeh et al. 2020). Based on the phylogenetic analysis including our samples and all three Iranian samples which are deposited in NCBI, this new subclade located as a sister subclade with *althaea*, with unresolved relationships between *halimodendri* and this new subclade (Fig. S3). Therefore, due to the lack of *minula COI* sequences in GenBank and the limited sampling from northeastern Iran, we propose that this new subclade should be considered a sister taxon to *halimodendri* and *althea*. However, further investigation is needed to determine whether this subclade represents a distinct population.

### Low intraspecific genetic distance and high morphological variation: a case study of Carduelis carduelis

The results revealed a split between two phenotypically different groups within *Carduelis carduelis*, the western *carduelis* group (black-headed subspecies group) and the eastern *caniceps* group (grey-headed subspecies group). However, this split was characterized by low genetic distance and low nodal support (Fig. 4, Table S5). In the *C. carduelis* complex, the *carduelis* subspecies group, ranges into the Zagros mountains in the west and north of Iran, whereas on the eastern side of its distributional range in Iran, it is replaced by the morphologically divergent *caniceps* group which its main range further north in Central Asia (Clements et al. 2023). These two subspecies groups indicate substantial differences in the plumage coloration and ornaments and there are some conflicts regarding their classification. E.g., Dickinson and Christidis (2014) consider the taxon *caniceps* as a subspecies of *C. carduelis* whereas, Gill and Donsker (2024) consider these two taxa as separate species i.e., *C. carduelis* and *C. caniceps* and, Clement et al. (2023) consider as two different subspecies groups. However, based on our results, these two taxa did not exhibit sufficient genetic differentiation to be considered separate species. Therefore, we propose that they be treated as two distinct subspecies groups, in accordance with their morphological variations, and different geographical distribution patterns. Nevertheless, it is suggested that ornaments may hinder gene flow between distinct or partially distinct populations if shaped by ecological differences or reinforcement, and this ability to modify ornaments, through mechanisms like sexual selection or reinforcement, could influence the formation of new species over time (Cardoso and Mota 2008).

### Effect of selection on COI gene

According to population genetic theory, the effectiveness of selection is more pronounced in species with larger effective population sizes (Woolfit 2009; Gossmann et al. 2010; Gossmann et al. 2012). Here we tested the effect of selection on protein coding mutations in the *COI* gene by contrasting the genetic diversity of mutations that change the amino acid at 0-fold degenerate sites versus those mutations that are silent at 4-fold degenerate sites (James et al. 2016). Species with larger effective population sizes should show relatively fewer amino-acid changing mutations because the selection is more efficient in larger populations. We were able to obtain non-zero coding diversity measures for 18 species and find a correlation between log (π 4-fold/0-fold) versus log (π 0-fold). The slope is negative and highly significant which suggests that much of the genetic variation is consistent with population size scaled effects of purifying selection and drift. Our results also suggest that much of the observed variation stems from mutations at synonymous sites.

### Conclusions

The patterns of intraspecific divergence observed in *S. tephronota*, *C. curruca* and *C. carduelis* align with key zoogeographic boundaries in Iran, reflecting an east-west geographical split within these species. This finding underscores the important role of DNA barcodes in revealing phylogeographical patterns, consistent with previous studies demonstrating the effectiveness of DNA barcoding in resolving such patterns (Saitoh et al. 2015; Wu et al. 2023). In addition, this east-west genetic distinctiveness in our target taxa is paralleled in several other widespread Eurasian passerine taxa that have allopatric populations at their southern range margin in Iran, such as the Eurasian nuthatch, *S. europaea* (Nazarizadeh et al. 2016; Päckert et al. 2020), coal tit, *Periparus ater* (Tietze et al. 2011), horned lark, *Eremophila alpestris* (Ghorbani et al. 2020) and great tit, *Parus major* (Javaheri Tehrani et al. 2021). These examples further highlight Iran’s pivotal role as a biogeography crossroad for avian diversity, with pair species consisting of a western (European or Mediterranean) member and an eastern (Asian) member (Haffer 1977; Roselaar and Aliabadian 2007). These findings collectively underscore the complex impact of Iran’s topography and climatic history on shaping present-day avian genetic diversity (Noori et al. 2024).

## Acknowledgments

This research was supported by Ferdowsi University of Mashhad research grant INSF 4002006 to Mansour Aliabadian and by a grant 3971 from Ferdowsi University of Mashhad awarded to Sahar Javaheri Tehrani. We gratefully acknowledge Sajad Noori for his assistance in creating the location map for our study and extend our thanks to Justin Wilcox for his support in enhancing the fluency of the English language in this manuscript. We also thank Iran’s Environmental Organization for their help during the field sampling and providing the necessary research permits for this study. We are particularly grateful to R. Rezaei for providing birds digital drawing. Anonymous reviewers are acknowledged, whose comments contributed to improvement of the manuscript.

## Data availability statement

Sequence data and associated collection information are will be available on the BOLD data system after acceptance and in the dataset entitled “DNA barcode reference library of Iranian passerine birds”.

## Author Contributions

**Investigation:** NAK, LN, VE, SK, MK, FA, SJT; **Resources:** NAK, LN, VE, SK, MK, FA, SJT, AS; **Formal analysis:** SJT, TIG, ER; **Writing – original draft:** SJT; **Funding acquisition:** MA, SJT; **Project administration:** MA; **Supervision:** TIG. MA; All authors read and approved the final version of the manuscript.

