## Supporting information for "DNA barcoding of passerine birds at an ornithological crossroad reveals significant East-West genetic lineage divergence"

**Supplementary Material (captions)**

**Figure S1.** Sampling locations. Colors and the size of circles represents the number of individuals.

**Figure S2.** Bayesian tree of *COI* sequences from 96 species of Iranian passerine birds. Values at nodes show posterior probabilities; full support is indicated with an asterisk. Species that form reciprocally monophyletic clades have been collapsed. Two species showed deep intraspecific divergence: (a) *Curruca curruca* and (b) *Sitta tephronota*; these are marked in blue. Families labeled on the right of the figure.

**Figure S3.** Phylogenetic and haplotype network analysis of *COI* data for *C. curruca*. a) Neighbor-joining tree, values on the branches shows bootstrap values and, our *COI* sequences are indicated in red. b) Haplotype network, where colors indicate the origin of the haplotypes (Orange: western population; Green: eastern populations) and the number of bars at each branch indicates the number of mutations.

**Table S1.** List of all Iranian passerine birds that have been sequenced in this study, with voucher numbers and collection localities. Coordinates are given in decimal degrees.

**Table S2.** Comparisons of K_2_P-pairwise distances within species. Distances are calculated for Iranian passerine birds for which two or more sequences were available; species including one individual are not calculated (n/c); three challenging taxa indicated bold and highlighted grey; distances are expressed in percentages.

**Table S3.** K2P distances (%) for the populations of *S. tephronota* and *S. neumayer* (*COI* seq), below the diagonal between group average.

**Table S4.** K2P distances (%) for the populations of *Curruca curruca* (*COI* seq), below the diagonal between group average.

**Table S5.** K2P distances (%) for the populations of *Carduelis carduelis* (*COI* seq), below the diagonal between group average.

**Table S6.** Genetic diversity at coding sites. Non-synonymous nucleotide diversity πn, number of non-synonymous nucleotide diversity N, synonymous nucleotide diversity πs, number of synonymous nucleotide diversity S.

**Supplementary Figures**

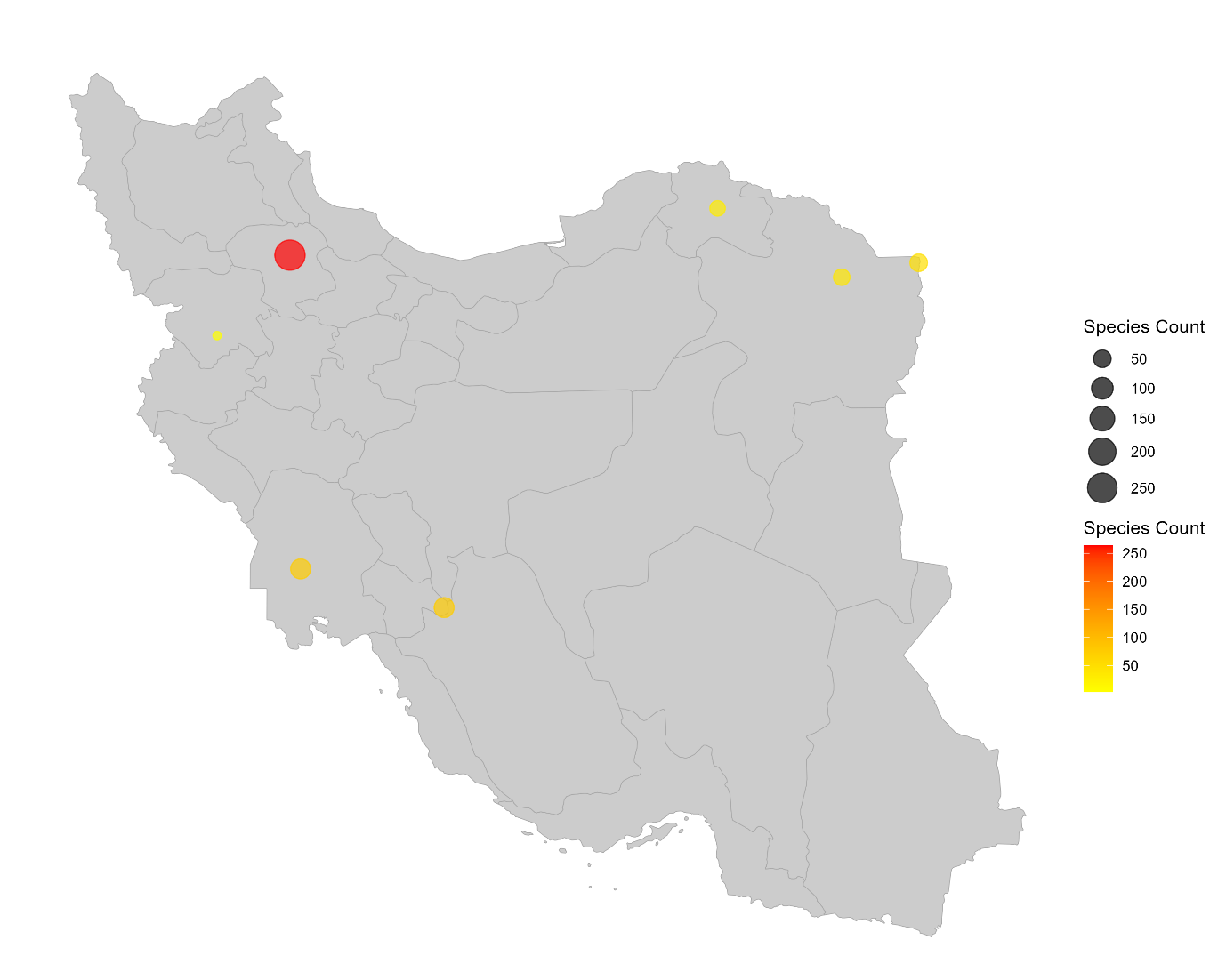

**Figure S1. Sampling locations.** Colors and the size of circles represents the number of individuals.

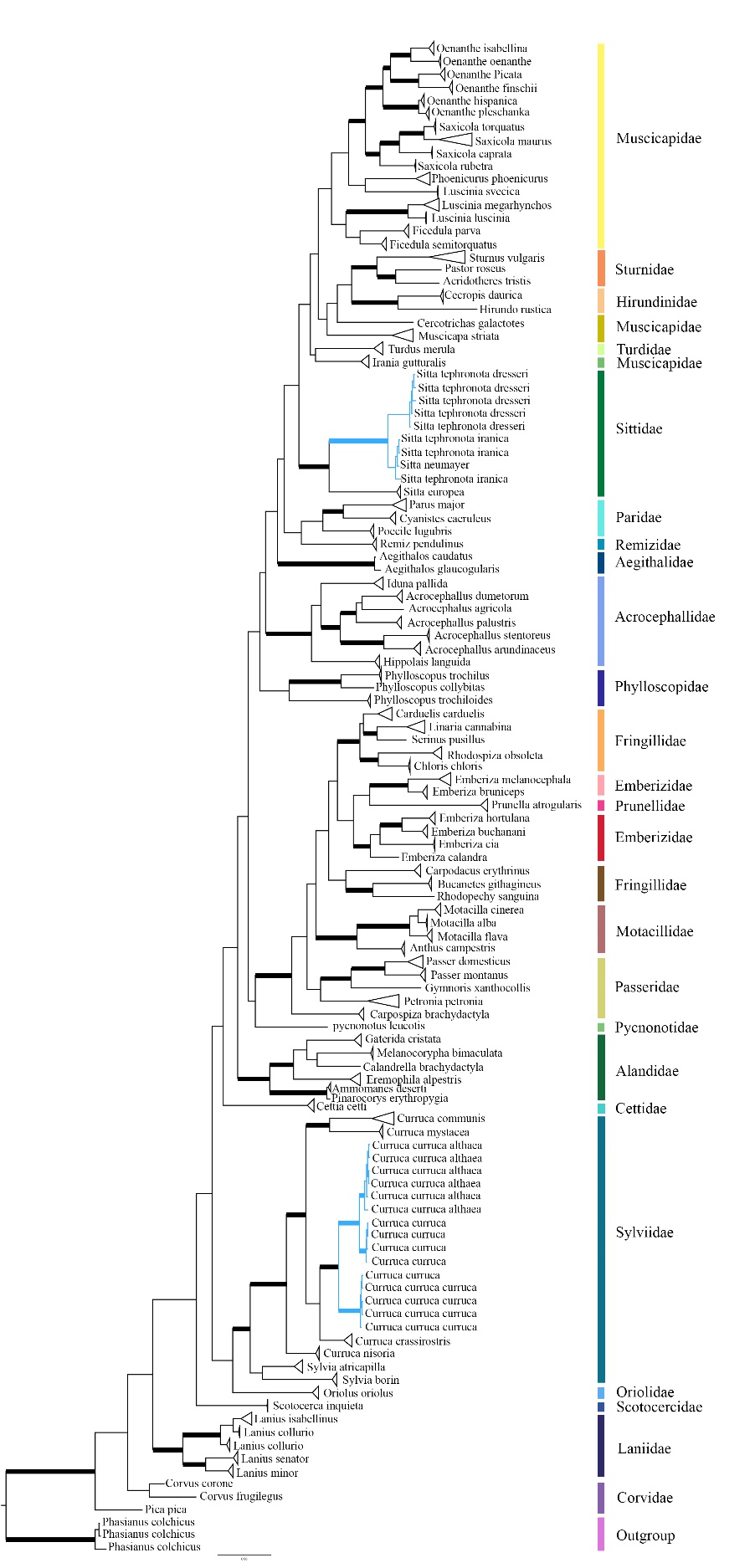

**Figure S2. Bayesian tree of *COI* sequences from 96 species of Iranian passerine birds.** Internal branches with posterior probabilities ≥ 0.95 indicates bold. Species that form reciprocally monophyletic clades have been collapsed. Two species showed deep intraspecific divergence: (a) *Curruca curruca* and (b) *Sitta tephronota*; these are marked in blue. Families labeled on the right of the figure.

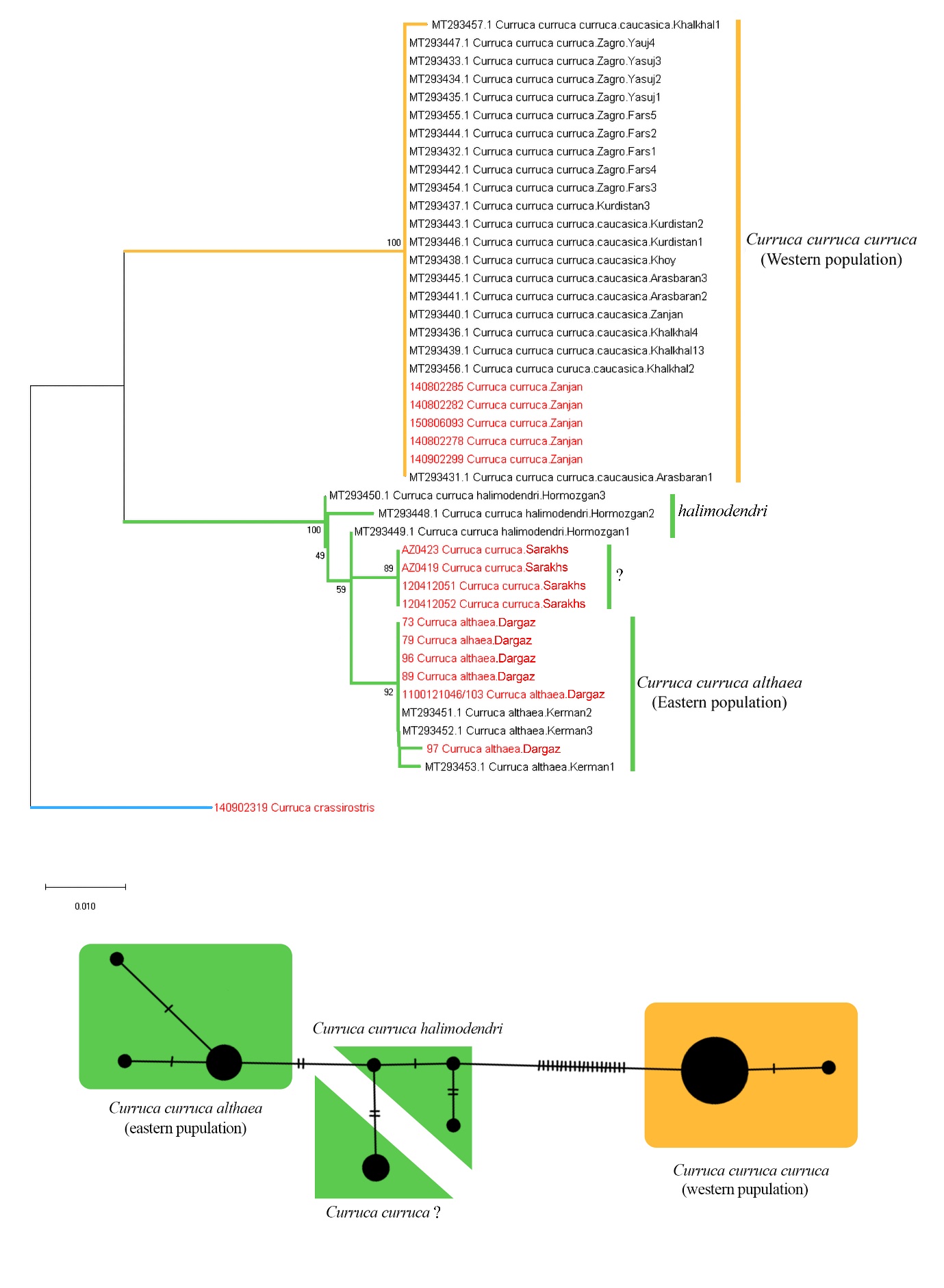

**Figure S3. Phylogenetic and haplotype network analysis of *COI* data for *C. curruca*.** **a)** Neighbor-joining tree, values on the branches shows bootstrap values and, our *COI* sequences are indicated in red. **b)** Haplotype network, where colors indicate the origin of the haplotypes (Orange: western population; Green: eastern populations) and the number of bars at each branch indicates the number of mutations.

**Supplementary Tables**

**Table S1.** **List of all Iranian passerine birds that have been sequenced in this study.** This table includes voucher numbers, localities and access numbers.

| **Species or subspecies** | **FUM number** | **Locality** | **Accession NO.** |
| --- | --- | --- | --- |
| *Luscinia luscinia* | 140802308 | Zanjan |  |
|  | 140902317 | Zanjan |  |
| *Luscinia megarhynchos* | 135 | Khorasan |  |
|  | 108 | Khorasan |  |
|  | 39 | Khorasan |  |
|  | 130706001 | Khorasan |  |
|  | S10 | Yasuj |  |
|  | 160516131 | Khuzestan |  |
|  | 160516129 | Khuzestan |  |
|  | 160416062 | Khuzestan |  |
|  | 130706070 | Zanjan |  |
|  | S01 | Yasuj |  |
| *Ficedula semitorquata* | 150806081 | Zanjan |  |
|  | 140802242 | Zanjan |  |
| *Ficedula parva* | 140802312 | Zanjan |  |
|  | 131 | Khorasan |  |
|  | AZ0421 | Khorasan |  |
|  | 120412056 | Sarakhs |  |
|  | 120412067 | Sarakhs |  |
|  | 120412059 | Sarakhs |  |
| *Oenanthe isabellina* | 130706063 | Zanjan |  |
|  | 130706065 | Zanjan |  |
|  | 150806110 | Zanjan |  |
|  | 1204061162 | Zanjan |  |
|  | 1204061161 | Zanjan |  |
|  | 140706080 | Zanjan |  |
|  | 150916213 | Khuzestan |  |
|  | 1204061126 | Zanjan |  |
| *Oenanthe oenanthe* | 1204061125 | Zanjan |  |
|  | 1204061140 | Zanjan |  |
|  | 1204061127 | Zanjan |  |
| *Oenanthe finschii* | 1204061138 | Zanjan |  |
|  | 160516146 | Khuzestan |  |
|  | 160516147 | Khuzestan |  |
|  | 160416117 | Khuzestan |  |
| *Oenanthe picata* | 120612117 | Sarakhs |  |
|  | 120712444 | Sarakhs |  |
|  | 70 | Khorasan |  |
|  | 95 | Khorasan |  |
|  | 76 | Khorasan |  |
|  | 80 | Khorasan |  |
| *Oenanthe hispanica* | 140706079 | Zanjan |  |
|  | 150806082 | Zanjan |  |
|  | 140802265 | Zanjan |  |
|  | 140802284 | Zanjan |  |
|  | 140706133 | Zanjan |  |
|  | 140802275 | Zanjan |  |
|  | 140802289 | Zanjan |  |
|  | 140802277 | Zanjan |  |
|  | 160416097 | Khuzestan |  |
| *Oenanthe pleschanka* | 140706097 | Zanjan |  |
|  | 140706041 | Zanjan |  |
|  | 140706054 | Zanjan |  |
| *Saxicola rubetra* | 160416110 | Khuzestan |  |
|  | 160416105 | Khuzestan |  |
| *Saxicola maurus* | 1204061121 | Zanjan |  |
|  | 1204061133 | Zanjan |  |
|  | 1204061135 | Zanjan |  |
|  | 41 | Khorasan |  |
|  | 44 | Khorasan |  |
| *Saxicola torquatus* | 140806141 | Zanjan |  |
| *Saxicola caprata* | 120412055 | Sarakhs |  |
|  | 120512093 | Sarakhs |  |
| *Phoenicurus phoenicurus* | 140902315 | Zanjan |  |
|  | k89 | Yasuj |  |
|  | 160416107 | Khuzestan |  |
|  | 160416088 | Khuzestan |  |
|  | k90 | Yasuj |  |
| *Irania_gutturalis* | 160416094 | Khuzestan |  |
|  | 160416108 | Khuzestan |  |
|  | 140806182 | Zanjan |  |
|  | 150806157 | Zanjan |  |
|  | 140706093 | Zanjan |  |
| *Luscinia svecica* | 150806132 | Zanjan |  |
|  | 150806136 | Zanjan |  |
| *Parus major* | 140706091 | Zanjan |  |
|  | 140802255 | Zanjan |  |
|  | 140902310 | Zanjan |  |
|  | 140802256 | Zanjan |  |
|  | 140802245 | Zanjan |  |
|  | 14 | Khorasan |  |
|  | 22 | Khorasan |  |
|  | 65 | Khorasan |  |
|  | K05 | Yasuj |  |
|  | K66 | Yasuj |  |
|  | K67 | Yasuj |  |
|  | S17 | Yasuj |  |
|  | S04 | Yasuj |  |
|  | K68 | Yasuj |  |
|  | K77 | Yasuj |  |
|  | S40 | Yasuj |  |
|  | S21 | Yasuj |  |
|  | S37 | Yasuj |  |
|  | 140802244 | Zanjan |  |
|  | 140706134d | Zanjan |  |
|  | 13 | Khorasan |  |
|  | 67 | Khorasan |  |
|  | 140802246 | Zanjan |  |
|  | 66 | Khorasan |  |
|  | 21 | Khorasan |  |
|  | S39 | Yasuj |  |
|  | 160416066 | Khuzestan |  |
|  | 150916202 | Khuzestan |  |
|  | S20 | Yasuj |  |
|  | S34 | Yasuj |  |
|  | S11 | Yasuj |  |
|  | S18 | Yasuj |  |
| *Poecile lugubris* | 1509162177 | Khuzestan |  |
|  | 160416109 | Khuzestan |  |
|  | 160416104 | Khuzestan |  |
|  | 150916206 | Khuzestan |  |
|  | Kk85 | Yasuj |  |
| *Cyanistes caeruleus* | 130722040 | Kurdistan |  |
|  | S15 | Yasuj |  |
|  | K78 | Yasuj |  |
|  | 160516134 | Khuzestan |  |
|  | 160516132 | Khuzestan |  |
|  | 150916210 | Khuzestan |  |
| Turdus merula | 160916066 | Khuzestan |  |
|  | K79 | Yasuj |  |
|  | S26 | Yasuj |  |
|  | S03 | Yasuj |  |
|  | 11 | Khorasan |  |
|  | 10 | Khorasan |  |
|  | S12 | Yasuj |  |
| *Muscicapa striata* | 140802241 | Zanjan |  |
|  | 140902301 | Zanjan |  |
|  | 150916204 | Khuzestan |  |
|  | 160516126 | Khuzestan |  |
|  | 140802287 | Zanjan |  |
|  | 120612126 | Sarakhs |  |
|  | 98 | Khorasan |  |
|  | 102 | Khorasan |  |
|  | 101 | Khorasan |  |
|  | 112 | Khorasan |  |
|  | 119 | Khorasan |  |
| *Remiz pendulinus* | 150806089 | Zanjan |  |
|  | 150806090 | Zanjan |  |
|  | 150806083 | Zanjan |  |
|  | 150806088 | Zanjan |  |
|  | 150806091 | Zanjan |  |
|  | 160916068 | Khuzestan |  |
| *Pycnonotus leucotis* | 150916218 | Khuzestan |  |
| *Curruca curruca* | 150806093 | Zanjan |  |
|  | 140802278 | Zanjan |  |
|  | 140802282 | Zanjan |  |
|  | 140902299 | Zanjan |  |
|  | 140802285 | Zanjan |  |
|  | 120412052 | Sarakhs |  |
|  | 120412051 | Sarakhs |  |
|  | AZ0423 | Sarakhs |  |
|  | AZ0419 | Sarakhs |  |
|  | 73 | Khorasan |  |
|  | 79 | Khorasan |  |
|  | 96 | Khorasan |  |
|  | 89 | Khorasan |  |
|  | 1100121046/103 | Khorasan |  |
|  | 97 | Khorasan |  |
| *Curruca crassirostris* | 140902319 | Zanjan |  |
|  | 140802268 | Zanjan |  |
|  | 140802288 | Zanjan |  |
|  | 140802290 | Zanjan |  |
|  | 150806092 | Zanjan |  |
| *Sylvia hortensis* | 120712438 | Sarakhs |  |
|  | 120712439 | Sarakhs |  |
|  | 120712440 | Sarakhs |  |
| *Curruca nisoria* | 140802261 | Zanjan |  |
|  | 140802267 | Zanjan |  |
|  | 140802271 | Zanjan |  |
|  | 140802262 | Zanjan |  |
| *Sylvia communis*  *Or Curruca communis* | 140706045 | Zanjan |  |
|  | 140706077 | Zanjan |  |
|  | 140706074 | Zanjan |  |
|  | 140706103 | Zanjan |  |
|  | 140706105 | Zanjan |  |
|  | 140902302 | Zanjan |  |
|  | 140706089 | Zanjan |  |
|  | 140706124 | Zanjan |  |
|  | 140706070 | Zanjan |  |
|  | 140802273 | Zanjan |  |
|  | 140802248 | Zanjan |  |
|  | K88 | Yasuj |  |
|  | 140706111 | Zanjan |  |
|  | 150806106 | Zanjan |  |
|  | 140802260 | Zanjan |  |
|  | 140706064 | Zanjan |  |
|  | 160916060 | Khuzestan |  |
|  | 160916061 | Khuzestan |  |
|  | 160516122 | Khuzestan |  |
|  | 150916211 | Khuzestan |  |
|  | 160916079 | Khuzestan |  |
|  | 140706138 | Zanjan |  |
|  | 140706130 | Zanjan |  |
|  | 140706108 | Zanjan |  |
|  | 140706047 | Zanjan |  |
|  | 140706053 | Zanjan |  |
|  | 140802269 | Zanjan |  |
|  | 140802270 | Zanjan |  |
|  | 140802272 | Zanjan |  |
|  | 150806099 | Zanjan |  |
|  | 56 | Khorasan |  |
|  | 38 | Khorasan |  |
| *Sylvia mystacea* | 150806064 | Zanjan |  |
|  | 150806077 | Zanjan |  |
|  | 150806120 | Zanjan |  |
|  | 150806085 | Zanjan |  |
| *Sylvia atricapilla* | 140802257 | Zanjan |  |
|  | 140902314 | Zanjan |  |
|  | 140902309 | Zanjan |  |
|  | S35 | Yasuj |  |
|  | S28 | Yasuj |  |
|  | S42 | Yasuj |  |
|  | 140902311 | Zanjan |  |
|  | 140902296 | Zanjan |  |
|  | 160916080 | Khuzestan |  |
|  | 160416078 | Khuzestan |  |
|  | 160416103 | Khuzestan |  |
|  | S41 | Yasuj |  |
| *Sylvia* *borin* | 140802253 | Zanjan |  |
|  | 140802254 | Zanjan |  |
|  | 140802258 | Zanjan |  |
|  | K75 | Yasuj |  |
|  | 140902306 | Zanjan |  |
|  | 160916075 | Khuzestan |  |
|  | 160916086 | Khuzestan |  |
| Oriolus oriolus | 140802251 | Zanjan |  |
|  | 140802249 | Zanjan |  |
|  | 140802266 | Zanjan |  |
|  | 140802250 | Zanjan |  |
| *Sturnus vulgaris* | 140706043 | Zanjan |  |
|  | 140706104 | Zanjan |  |
|  | 150806068 | Zanjan |  |
|  | 150806069 | Zanjan |  |
|  | 1204061147 | Zanjan |  |
|  | CH3214 | Sarakhs |  |
|  | CH3216 | Sarakhs |  |
|  | CH3217 | Sarakhs |  |
|  | 150806070 | Zanjan |  |
|  | 1204061149 | Zanjan |  |
|  | 130 | Khorasan |  |
|  | 140706095 | Zanjan |  |
| *Pastor roseus* | 140802237 | Zanjan |  |
| *Acridotheres tristis* | 94 | Khorasan |  |
| *Sitta tephronata* | 140806192 | Zanjan |  |
|  | 150916200 | Khuzestan |  |
|  | 160416100 | Khuzestan |  |
|  | 160416102 | Khuzestan |  |
|  | 140902321 | Zanjan |  |
|  | 120612136 | Sarakhs |  |
|  | 120612124 | Sarakhs |  |
| *Sitta tephronata iranica* | 133 | Khorasan |  |
|  | 111 | Khorasan |  |
| *Sitta europaea* | 130722027 | Kurdstan |  |
|  | K83 | Yasuj |  |
|  | K87 | Yasuj |  |
| *Lanius senator* | 130706058 | Zanjan |  |
|  | 140802264 | Zanjan |  |
|  | 140802281 | Zanjan |  |
|  | 140802280 | Zanjan |  |
|  | 160416067 | Khuzestan |  |
|  | 150806163 | Zanjan |  |
| *Lanius minor* | 130706071 | Zanjan |  |
|  | 1204061157 | Zanjan |  |
|  | 140706066 | Zanjan |  |
| *Lanius isabellinus* | S36 | Yasuj |  |
|  | 120712437 | Sarakhs |  |
| *Lanius collurio* | 140802247 | Zanjan |  |
|  | 140902305 | Zanjan |  |
|  | 140802274 | Zanjan |  |
|  | S27 | Yasuj |  |
|  | K64 | Yasuj |  |
|  | K50 | Yasuj |  |
|  | 140902298 | Zanjan |  |
|  | 140902294 | Zanjan |  |
|  | 130706059 | Zanjan |  |
|  | 140802286 | Zanjan |  |
| *corvus corone* | K80 | Yasuj |  |
| *Corvus frugilegus* | 125 | Khorasan |  |
| *Pica pica* | 139 | Khorasan |  |
| *Phylloscopus collybitas* | 1204061150 | Zanjan |  |
| *Phylloscopus trochilus* | 160916072 | Khuzestan |  |
|  | S30 | Yasuj |  |
|  | S31 | Yasuj |  |
| *Scotocerca inquieta* | 160416056 | Khuzestan |  |
|  | 160416073 | Khuzestan |  |
| *Phyloscopus trochiloides* | 120412057 | Sarakhs |  |
|  | 120512109 | Sarakhs |  |
| *Carduelis carduelis* | 140806156 | Zanjan |  |
|  | 140802238 | Zanjan |  |
|  | 140802283 | Zanjan |  |
|  | 140802279 | Zanjan |  |
|  | S09 | Yasuj |  |
|  | 160416070 | Khuzestan |  |
|  | 160416068 | Khuzestan |  |
|  | 160416071 | Khuzestan |  |
|  | S33 | Yasuj |  |
|  | 120512082 | Sarakhs |  |
|  | 120612138 | Sarakhs |  |
|  | 140706075 | Zanjan |  |
|  | 128 |  |  |
|  | 1408061566 | Zanjan |  |
|  | 122 | Khorasan |  |
| *Linaria cannabina* | 140806209 | Zanjan |  |
|  | 140806197 | Zanjan |  |
|  | 150806146 | Zanjan |  |
|  | 1204061152 | Zanjan |  |
|  | 160916087 | Khuzestan |  |
|  | 160916076 | Khuzestan |  |
|  | 140806202 | Zanjan |  |
|  | 140806204 | Zanjan |  |
|  | 140806206 | Zanjan |  |
|  | 140706115 | Zanjan |  |
|  | 140806199 | Zanjan |  |
|  | 136 | Sarakhs |  |
|  | 130706064 | Zanjan |  |
| *Serinus pusillus* | K102 | Yasuj |  |
| *Rhodospiza obsoleta* | 140706048 | Zanjan |  |
|  | 140706055 | Zanjan |  |
|  | 120612123 | Sarakhs |  |
|  | 160516145 | Yasuj |  |
|  | K95 | Yasuj |  |
|  | 160416089 | Khuzestan |  |
|  | 137 | Sarakhs |  |
|  | 138 | Sarakhs |  |
|  | 26 | Khorasan |  |
| *Chloris chloris* | 42 | Khorasan |  |
|  | 46 | Khorasan |  |
| *Emberiza calandra* | 130706066 | Zanjan |  |
| *Emberiza buchanani* | 140806189 | Zanjan |  |
|  | 140706119 | Zanjan |  |
|  | 150806118 | Zanjan |  |
|  | 140706118 | Zanjan |  |
| *Emberiza hortulana* | 140802276 | Khuzestan |  |
|  | 140902320 | Zanjan |  |
|  | 140902291 | Zanjan |  |
|  | 120512101 | Sarakhs |  |
| *Emberiza cia* | 140802263 | Zanjan |  |
|  | 140SCOI | Sarakhs |  |
| *Emberiza melanocephala* | 140706039 | Zanjan |  |
|  | 140706128 | Zanjan |  |
|  | 1204061123 | Zanjan |  |
|  | 130706060 | Zanjan |  |
|  | 160416114 | Khuzestan |  |
|  | 160416076 | Khuzestan |  |
|  | 150806049 | Zanjan |  |
|  | 130722089 | Kurdstan |  |
|  | 1204061122 | Zanjan |  |
|  | 1204061136 | Zanjan |  |
|  | 160416079 | Khuzestan |  |
| *Emberiza bruniceps* | 1205121117 | Sarakhs |  |
|  | 48 | Khorasan |  |
|  | 92 | Khorasan |  |
|  | 78 | Khorasan |  |
| *Rhodopechys sanguina* | 150806114 | Zanjan |  |
| *Bucanetes githagineus* | 120612133 | Sarakhs |  |
|  | 120612134 | Sarakhs |  |
| *Motacilla flava* | 120406120 | Zanjan |  |
|  | 1204061131 | Zanjan |  |
| *Motacilla alba* | 124 | Khorasan |  |
|  | 120812629 | Sarakhs |  |
| *Motacilla cinerea* | S08 | Yasuj |  |
|  | K101 | Yasuj |  |
|  | 150916212 | Khuzestan |  |
|  | 160916067 | Khuzestan |  |
| *Anthus campestris* | 1204061145 | Zanjan |  |
|  | 1204061146 | Zanjan |  |
|  | 1204061154 | Zanjan |  |
| *Petronia petronia* | 140706029 | Zanjan |  |
|  | 140706035 | Zanjan |  |
|  | 140706072 | Zanjan |  |
|  | 120512078 | Sarakhs |  |
|  | 120612131 | Sarakhs |  |
|  | K98 | Yasuj |  |
|  | K99 | Yasuj |  |
|  | 140706078 | Zanjan |  |
|  | 144 | Khorasan |  |
|  | 82 | Khorasan |  |
|  | 81 | Khorasan |  |
|  | 140706065 | Zanjan |  |
|  | 140706033 | Zanjan |  |
|  | 140706058 | Zanjan |  |
|  | 140706051 | Zanjan |  |
|  | 140706057 | Zanjan |  |
| *Carpodacus erythrinus* | 140802252 | Zanjan |  |
|  | 140802259 | Zanjan |  |
|  | 140902316 | Zanjan |  |
|  | 140902313 | Zanjan |  |
|  | 140902318 | Zanjan |  |
| *Passer domesticus* | 140806196 | Zanjan |  |
|  | 140802243 | Zanjan |  |
|  | 31 | Khorasan |  |
|  | K61 | Yasuj |  |
|  | K74 | Yasuj |  |
|  | 120412066 | Sarakhs |  |
|  | 120412062 | Sarakhs |  |
|  | 160916078 | Khuzestan |  |
|  | 160916083 | Khuzestan |  |
|  | 160916073 | Khuzestan |  |
|  | 130706009 | Zanjan |  |
|  | 140902303 | Zanjan |  |
|  | 140902307 | Zanjan |  |
|  | S29 | Yasuj |  |
|  | K63 | Yasuj |  |
|  | K55 | Yasuj |  |
|  | 160916081 | Khuzestan |  |
|  | 160916085 | Khuzestan |  |
|  | 140802239 | Zanjan |  |
|  | 120412063 | Sarakhs |  |
|  | 130706008 | Zanjan |  |
|  | 150806053 | Zanjan |  |
|  | 160416057 | Khuzestan |  |
|  | K51 | Yasuj |  |
|  | K62 | Yasuj |  |
|  | S16 | Yasuj |  |
|  | 130706006 | Zanjan |  |
|  | 28 | Khorasan |  |
|  | 34 | Khorasan |  |
|  | 50 | Khorasan |  |
| *Passer montanus* | 1204061167 | Zanjan |  |
|  | 120812920 | Sarakhs |  |
|  | 68 | Khorasan |  |
|  | 63 | Khorasan |  |
|  | 20 | Khorasan |  |
| *Gymnoris xanthocollis* | K94 | Yasuj |  |
| *Carpospiza brachydactyla* | 130706068 | Zanjan |  |
|  | 140706050 | Zanjan |  |
|  | 130706005 | Zanjan |  |
|  | 140706082 | Zanjan |  |
| *Prunella atrogularis* | 150806121 | Zanjan |  |
|  | 150806134 | Zanjan |  |
|  | 150806156 | Zanjan |  |
|  | 150806153 | Zanjan |  |
|  | 150806135 | Zanjan |  |
|  | 150806137 | Zanjan |  |
|  | 150806140 | Zanjan |  |
|  | 150806155 | Zanjan |  |
|  | 150806154 | Zanjan |  |
| *Cettia cetti* | 140706062 | Zanjan |  |
|  | 140706084 | Zanjan |  |
|  | 140706131 | Zanjan |  |
|  | 150806123 | Zanjan |  |
|  | 150806102 | Zanjan |  |
|  | K56 | Yasuj |  |
|  | K59 | Yasuj |  |
|  | K72 | Yasuj |  |
|  | 160916062 | Khuzestan |  |
|  | K73 | Yasuj |  |
|  | 140706073 | Zanjan |  |
|  | 150806138 | Zanjan |  |
|  | 140706068 | Zanjan |  |
| *Galerida cristata* | 140706028 | Zanjan |  |
|  | 140706125 | Zanjan |  |
|  | 1204061156 | Zanjan |  |
|  | 140706100 | Zanjan |  |
|  | 1204061141 | Zanjan |  |
|  | 140902292 | Zanjan |  |
|  | 140902304 | Zanjan |  |
|  | 1204061155 | Zanjan |  |
|  | 140902293 | Zanjan |  |
|  | 1204061153 | Zanjan |  |
|  | 120812923 | Sarakhs |  |
|  | 120812921 | Sarakhs |  |
|  | 120812924 | Sarakhs |  |
|  | 120812928 | Sarakhs |  |
|  | 150916219 | Khuzestan |  |
|  | 150916217 | Khuzestan |  |
|  | 33 | Khorasan |  |
| *Melanocorypha bimaculata* | 1204061148 | Zanjan |  |
|  | 1204061169 | Zanjan |  |
|  | 1204061171 | Zanjan |  |
| *Calandrella brachydactyla* | 1204061165 | Zanjan |  |
| *Eremophila alpestris* | 150806117 | Zanjan |  |
|  | 1204061158 | Zanjan |  |
|  | 1204061144 | Zanjan |  |
|  | 150806144 | Zanjan |  |
|  | 120612135 | Sarakhs |  |
|  | 150806149 | Zanjan |  |
|  | 150806150 | Zanjan |  |
|  | 107 | Khorasan |  |
| *Pinarocorys erythropygia* | 140706110 | Zanjan |  |
| *Ammomanes deserti* | 160516144 | Khuzestan |  |
|  | 160516141 | Khuzestan |  |
| *Iduna pallida* | 140806151 | Zanjan |  |
|  | 150806072 | Zanjan |  |
|  | 150806074 | Zanjan |  |
|  | 140706099 | Zanjan |  |
|  | 150806066 | Zanjan |  |
|  | 140806088 | Zanjan |  |
|  | K52 | Yasuj |  |
|  | K93 | Yasuj |  |
|  | K53 | Yasuj |  |
|  | 140706076 | Zanjan |  |
|  | 140706060 | Zanjan |  |
|  | 140806161 | Zanjan |  |
|  | 1605161411 | Khuzestan |  |
|  | 160516136 | Khuzestan |  |
|  | 150806075 | Zanjan |  |
|  | K60 | Yasuj |  |
|  | K57 | Yasuj |  |
|  | 150806079 | Zanjan |  |
|  | 140706063 | Zanjan |  |
|  | 140706086 | Zanjan |  |
|  | 140806149 | Zanjan |  |
|  | K69 | Yasuj |  |
|  | 150806084 | Zanjan |  |
|  | 140806142 | Zanjan |  |
|  | K76 | Yasuj |  |
| *Acrocephallus palustris* | 1408061499 | Zanjan |  |
|  | K71 | Yasuj |  |
|  | K70 | Yasuj |  |
|  | 150806051_ | Zanjan |  |
|  | 140902295 | Zanjan |  |
|  | 160516128 | Khuzestan |  |
|  | 150806104 | Zanjan |  |
| *Acrocephalus dumetorum* | 120512110 | Sarakhs |  |
|  | 120512111 | Sarakhs |  |
|  | 120412053 | Sarakhs |  |
|  | 45 | Khorasan |  |
|  | 16 | Khorasan |  |
|  | 51 | Khorasan |  |
|  | 18 | Khorasan |  |
| *Acrocephallus agricola* | 116 | Khorasan |  |
| *Acrocephalus arundinaceus* | 130706002 | Zanjan |  |
|  | K65 | Yasuj |  |
| *Acrocephallus stentoreus* | 146 | Khorasan |  |
|  | 147 | Khorasan |  |
|  | 148 | Khorasan |  |
| *Hippolais languida* | 140706092 | Zanjan |  |
|  | 140706107 | Zanjan |  |
|  | 160516123 | Khuzestan |  |
|  | 160516127 | Khuzestan |  |
| *Cecropis daurica* | 160416119 | Khuzestan |  |
|  | 160416059 | Khuzestan |  |
|  | 160416060 | Khuzestan |  |
| *Hirundo rustica* | S25 | Yasuj |  |
| *Aegithalos caudatus* | 150916216 | Khuzestan |  |
| *Aegithalos glaucogularis vinaceus* | K96 | Yasuj |  |
| *Cercotrichas galactotes* | 17 | Khorasan |  |

**Table S2.** **Comparisons of K_2_P-pairwise distances within species.** Distances are calculated for Iranian passerine birds for which two or more sequences were available; species including one individual are not calculated (n/c); three challenging taxa indicated bold and highlighted grey; distances are expressed in percentages.

| **Species** | **Intraspecific K_2_P genetic distances** | **S.E** |
| --- | --- | --- |
| *Luscinia luscinia* | 0.00 | 0.0000 |
| *Luscinia megarhynchos* | 0.83 | 0.0025 |
| *Luscinia svecica* | 0.00 | 0.0000 |
| *Ficedula semitorquata* | 0.73 | 0.0029 |
| *Ficedula parva* | 0.13 | 0.0008 |
| *Oenanthe isabellina* | 0.10 | 0.0010 |
| *Oenanthe oenanthe* | 0.12 | 0.0011 |
| *Oenanthe finschii* | 0.12 | 0.0012 |
| *Oenanthe picata* | 0.32 | 0.0009 |
| *Oenanthe hispanica* | 0.00 | 0.0000 |
| *Oenanthe pleschanka* | 0.37 | 0.0021 |
| *Saxicola rubetra* | 0.00 | 0.0000 |
| *Saxicola torquatus* | 1.74 | 0.0041 |
| *Saxicola maurus* | 1.87 | 0.0064 |
| *Saxicola caprata* | 0.00 | 0.0000 |
| *Phoenicurus phoenicurus* | 1.09 | 0.0027 |
| *Irania gutturalis* | 0.48 | 0.0018 |
| *Muscicapa striata* | 1.08 | 0.0034 |
| *Cercotrichas galactotes* | n/c | n/c |
| *Turdus merula* | 0.42 | 0.0018 |
| *Parus major* | 0.04 | 0.0003 |
| *Cyanistes caeruleus* | 0.18 | 0.0011 |
| *Poecile lugubris* | 0.11 | 0.0012 |
| *Remiz pendulinus* | 0.00 | 0.0000 |
| *Pycnonotus leucotis* | n/c | n/c |
| ***Curruca curruca*** | **3.61** | **0.0067** |
| *Curruca crassirostris* | 0.22 | 0.0014 |
| *Curruca nisoria* | 0.09 | 0.0010 |
| *Curruca hortensis* | 0.12 | 0.0012 |
| *Curruca communis* | 0.47 | 0.0011 |
| *Curruca mystacea* | 0.00 | 0.0000 |
| *Sylvia atricapilla* | 0.17 | 0.0010 |
| *Sylvia borin* | 0.11 | 0.0011 |
| *Oriolus oriolus* | 0.40 | 0.0019 |
| *Galerida cristata* | 0.06 | 0.0004 |
| *Melanocorypha bimaculata* | 0.12 | 0.0013 |
| *Eremophila alpestris* | 0.44 | 0.0018 |
| *Calandrella brachydactyla* | 0.12 | 0.0013 |
| *Pinarocorys erythropygia* | n/c | n/c |
| *Ammomanes deserti* | 0.00 | 0.0000 |
| ***Sitta tephronata*** | **2.30** | **0.0048** |
| *Sitta neumayer* | n/c | n/c |
| *Sitta europaea* | 0.26 | 0.0016 |
| *Iduna pallida* | 0.28 | 0.0008 |
| *Hippolais languida* | 0.18 | 0.0013 |
| *Acrocephalus palustris* | 0.31 | 0.0012 |
| *Acrocephalus agricola* | n/c | n/c |
| *Acrocephalus dumetorum* | 0.12 | 0.0008 |
| *Acrocephalus arundinaceus* | 1.29 | 0.0043 |
| *Acrocephalus stentoreus* | 0.16 | 0.0018 |
| *Sturnus vulgaris* | 1.30 | 0.0025 |
| *Pastor roseus* | n/c | n/c |
| *Acridotheres tristis* | n/c | n/c |
| *Lanius senator* | 0.16 | 0.0011 |
| *Lanius minor* | 0.38 | 0.0022 |
| *Lanius isabellinus* | 0.00 | 0.0000 |
| *Lanius collurio* | 1.56 | 0.0034 |
| *Pica pica* | n/c | n/c |
| *Corvus corone* | n/c | n/c |
| *Corvus frugilegus* | n/c | n/c |
| *Cettia cetti* | 0.11 | 0.0006 |
| *Phylloscopus collybita* | n/c | n/c |
| *Phylloscopus trochiloides* | 0.36 | 0.0024 |
| *Phylloscopus trochilus* | 0.00 | 0.0000 |
| *Scotocerca inquieta* | 0.00 | 0.0000 |
| ***Carduelis carduelis*** | **0.43** | **0.0015** |
| *Linaria cannabina* | 0.65 | 0.0014 |
| *Serinus pusillus* | n/c | n/c |
| *Rhodospiza obsoleta* | 0.24 | 0.0014 |
| *Chloris chloris* | 0.00 | 0.0000 |
| *Carpodacus erythrinus* | 0.22 | 0.0012 |
| *Rhodopechys sanguineus* | n/c | n/c |
| *Bucanetes githagineus* | 0.36 | 0.0030 |
| *Passer domesticus* | 0.41 | 0.0009 |
| *Passer montanus* | 0.12 | 0.0015 |
| *Gymnoris xanthocollis* | 0.12 | 0.0015 |
| *Carpospiza brachydactyla* | 0.36 | 0.0021 |
| *Petronia petronia* | 1.46 | 0.0023 |
| *Motacilla flava* | 0.55 | 0.0035 |
| *Motacilla alba* | 0.55 | 0.0035 |
| *Motacilla cinerea* | 0.36 | 0.0018 |
| *Anthus campestris* | 0.12 | 0.0011 |
| *Prunella atrogularis* | 0.23 | 0.0010 |
| *Aegithalos caudatus* | n/c | n/c |
| *Aegithalos glaucogularis vinaceus* | n/c | n/c |
| *Cecropis daurica* | 0.00 | 0.0000 |
| *Hirundo rustica* | n/c | n/c |
| *Emberiza calandra* | n/c | n/c |
| *Emberiza buchanani* | 0.27 | 0.0019 |
| *Emberiza hortulana* | 0.18 | 0.0011 |
| *Emberiza cia* | 0.18 | 0.0016 |
| *Emberiza melanocephala* | 0.51 | 0.0013 |
| *Emberiza bruniceps* | 0.25 | 0.0018 |

**Table S3.** **K2P distances (%) for the populations of *S. tephronota* and *S. neumayer* (*COI* seq), below the diagonal between group average.**

| **Subspecies** | ***S. t. iranica*** | ***S. t. dresseri*** | ***S. neumayer***  **(pure)** |
| --- | --- | --- | --- |
| ***S. t. iranica*** |  |  |  |
| ***S. t. dresseri*** | 4.105 |  |  |
| ***S. neumayer* (pure)** | 5.933 | 6.451 |  |
| ***S. neumayer* (probable hybrid)** | 0.447 | 3.624 | 6.429 |

**Table S4.** **K2P distances (%) for the populations of *Curruca curruca* (*COI* seq), below the diagonal between group average.**

| **Subspecies** | ***C. curruca ?* (east)** | ***C. c. curuca* (west)** |
| --- | --- | --- |
| ***C. curruca ? (east)*** |  |  |
| ***C. c. curuca (west)*** | 6.99 |  |
| ***C. c. althaea (east)*** | 1.20 | 7.04 |

**Table S5. K2P distances (%) for the populations of *Carduelis carduelis* (*COI* seq), below the diagonal between group average.**

| **Subspecies** | ***caniceps* group** |
| --- | --- |
| ***caniceps* group** |  |
| ***carduelis* group** | 0.6671 |

**Table S6. Genetic diversity at coding sites.** Non-synonymous nucleotide diversity πn, number of non-synonymous nucleotide diversity N, synonymous nucleotide diversity πs, number of synonymous nucleotide diversity S.

| **Species** | **πn** | **N** | **πs** | **S** | **Log (πs)** | **Log (πn/πs)** |
| --- | --- | --- | --- | --- | --- | --- |
| *Carduelis cannabina* | 0.00211715 | 218 | 0.01118881 | 55 | -3.6915787 | -1.3211263 |
| *Carduelis carduelis* | 0.00108844 | 245 | 0.00689655 | 58 | -3.924796 | -1.4275674 |
| *Carpospiza brachydactyla* | 0.00148368 | 337 | 0.00694444 | 72 | -4.015695 | -1.3405948 |
| *Curruca communis* | 0.00067204 | 186 | 0.01703423 | 49 | -3.4588735 | -1.9832424 |
| *Emberiza melanocephala* | 0.00114351 | 318 | 0.01038961 | 70 | -3.8284988 | -1.6156882 |
| *Eremophila alpestris* | 0.00085034 | 294 | 0.01545842 | 67 | -3.6369096 | -1.901845 |
| *Erythrina erythrina* | 0.00118694 | 337 | 0.00555556 | 72 | -4.112605 | -1.3405948 |
| *Ficedula parva* | 0.00106838 | 312 | 0.00483092 | 69 | -4.1548194 | -1.310611 |
| *Ficedula semitorquata* | 0.00890208 | 337 | 0.01388889 | 72 | -3.714665 | -0.8634736 |
| *Hippolais pallida* | 0.00166172 | 337 | 0.00212963 | 72 | -4.5290284 | -0.7780434 |
| *Lanius collurio* | 0.00178042 | 337 | 0.0404321 | 72 | -3.2506062 | -2.0265023 |
| *Luscinia megarhynchos* | 0.00085837 | 233 | 0.02528736 | 58 | -3.3605246 | -2.0731573 |
| *Passer domesticus* | 0.00144928 | 184 | 0.01706258 | 45 | -3.4211678 | -1.6824991 |
| *Petronia petronia* | 0.01104827 | 221 | 0.04985119 | 56 | -3.0505125 | -1.2505857 |
| *Phoenicurus phoenicurus* | 0.00382166 | 314 | 0.03188406 | 69 | -3.3352755 | -1.5794025 |
| *Prunella atrogularis* | 0.00197824 | 337 | 0.00848765 | 72 | -3.9285448 | -1.3028062 |
| *Rhodopechys githaginea* | 0.00296736 | 337 | 0.02777778 | 72 | -3.413635 | -1.6416248 |
| *Sturnus vulgaris* | 0.01211015 | 274 | 0.02249053 | 64 | -3.4541803 | -0.9004207 |
